# Smart3-ATAC: a highly sensitive method for joint accessibility and full-length transcriptome analysis in single cells

**DOI:** 10.1101/2021.12.02.470912

**Authors:** Huaitao Cheng, Han-pin Pui, Antonio Lentini, Solrún Kolbeinsdóttir, Nathanael Andrews, Yu Pei, Björn Reinius, Qiaolin Deng, Martin Enge

## Abstract

Joint single-cell measurements of gene expression and DNA regulatory element activity holds great promise as a tool to understand transcriptional regulation. Towards this goal we have developed Smart3-ATAC, a highly sensitive method which allows joint mRNA and chromatin accessibility analysis genome wide in single cells. With Smart3-ATAC, we are able to obtain the highest possible quality measurements per cell. The method combines transcriptomic profiling based on the highly sensitive Smart-seq3 protocol on cytosolic mRNA, with a novel low-loss single-cell ATAC (scATAC) protocol to measure chromatin accessibility. Compared to current droplet multiome methods, the yield of both the scATAC protocol and mRNA-seq protocol is markedly higher.

## Background

Analysis of single-cell transcriptomes at a large scale is now routinely done, and has already produced highly detailed maps of cell states in health and disease. However, much of this data is descriptive in nature. Cell state is determined by the activity of epigenetic factors, such as transcription factors (TFs) acting on chromatin in a sequence specific manner. To understand the biology of the organism we therefore need to understand the rules that govern gene regulation. Joint measurements of gene expression and gene enhancer activity allow us to study correlations between the two, enabling an unbiased way to create hypotheses about the underlying biology of gene regulation (Lee, Hyeon, and Hwang 2020; S. Chen, Lake, and Zhang 2019). Previous approaches have focused on achieving high throughput measurements (Cao et al. 2018), but many scientific questions are better addressed by focusing on high precision and sensitivity rather than breadth. We therefore propose Smart3-ATAC - a highly sensitive method of joint chromatin accessibility and mRNA quantification which maximizes the information content of each single cell, while still maintaining throughput and cost effectiveness. The scATAC data is of comparable specificity to current gold standard methods, but with larger library sizes while the mRNA-seq is based on Smart-seq3, achieving highlly sensitive full-length transcript read coverage and an mechanism for counting original mRNA molecules via 5’ encoded unique molecular identifiers (UMIs).

### Development of a high sensitivity method for joint accessibility and mRNA profiling in single cells

Smart3-ATAC was specifically developed to obtain high sensitivity and information content measurements of both chromatin accessibility and mRNA expression. In order to achieve high precision measurements of the transcriptome, it is necessary to analyze cytosolic mRNA. Cytosolic and nuclear fractions from a single cell can be readily separated using centrifugation followed by automated pipetting in microwells. The nuclear chromatin can subsequently be tagmented using tn5 loaded with adapters, and barcoded in a similar manner to whole genome sequencing in DNTR-seq (Zachariadis et al. 2020). However, this leads to a technical problem since efficient tagmentation requires high concentration of transposome, but the resulting library sizes are so small that PCR amplification will fail unless the excess tn5 adaptor sequences are removed. Previous scATAC-seq methods have solved this problem by performing tagmentation on bulk samples of nuclei where the transposome can be easily washed away before separating into single nuclei(X. Chen et al. 2018; Cusanovich et al. 2015). However, this precludes the possibility to analyze cytosolic mRNA. In Smart3-ATAC, illustrated in Fig. 1, we instead sort single intact cells into nucleus extraction buffer, then mechanically separate the cytosolic fraction from the nucleus which we then tagment and wash individually in its well prior to barcoding PCR. By washing the intact nucleus we achieve a buffer exchange to remove free adapters, while avoiding a lossy DNA clean-up step.

**Figure 1.**
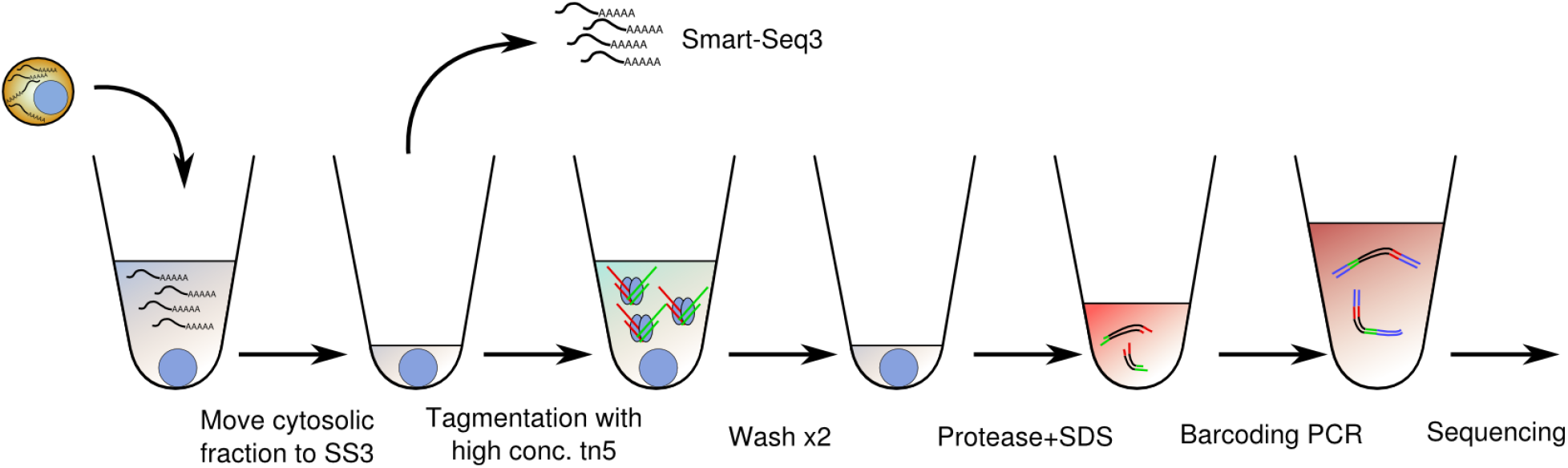
Summary of the method.

### Application of Smart3-ATAC to study *in vitro* gastrulation

To test the performance of Smart3-ATAC on a complex cell differentiation process, we analyzed 3196 cells harvested at four different time-points (0hr, 72hr, 96hr and 120hr) using stem cell-based *in vitro* modeling of mouse gastrulation(Beccari et al. 2018). First, we analyzed the quality of the resulting joint scATAC and mRNA-seq libraries. The scATAC libraries had a similar specificity to current-gen scATAC but with higher yield (Fig 2 a-d), averaging 38 417 reads per cell compared to 12 225 for 10x genomics multiome.

**Figure 2.**
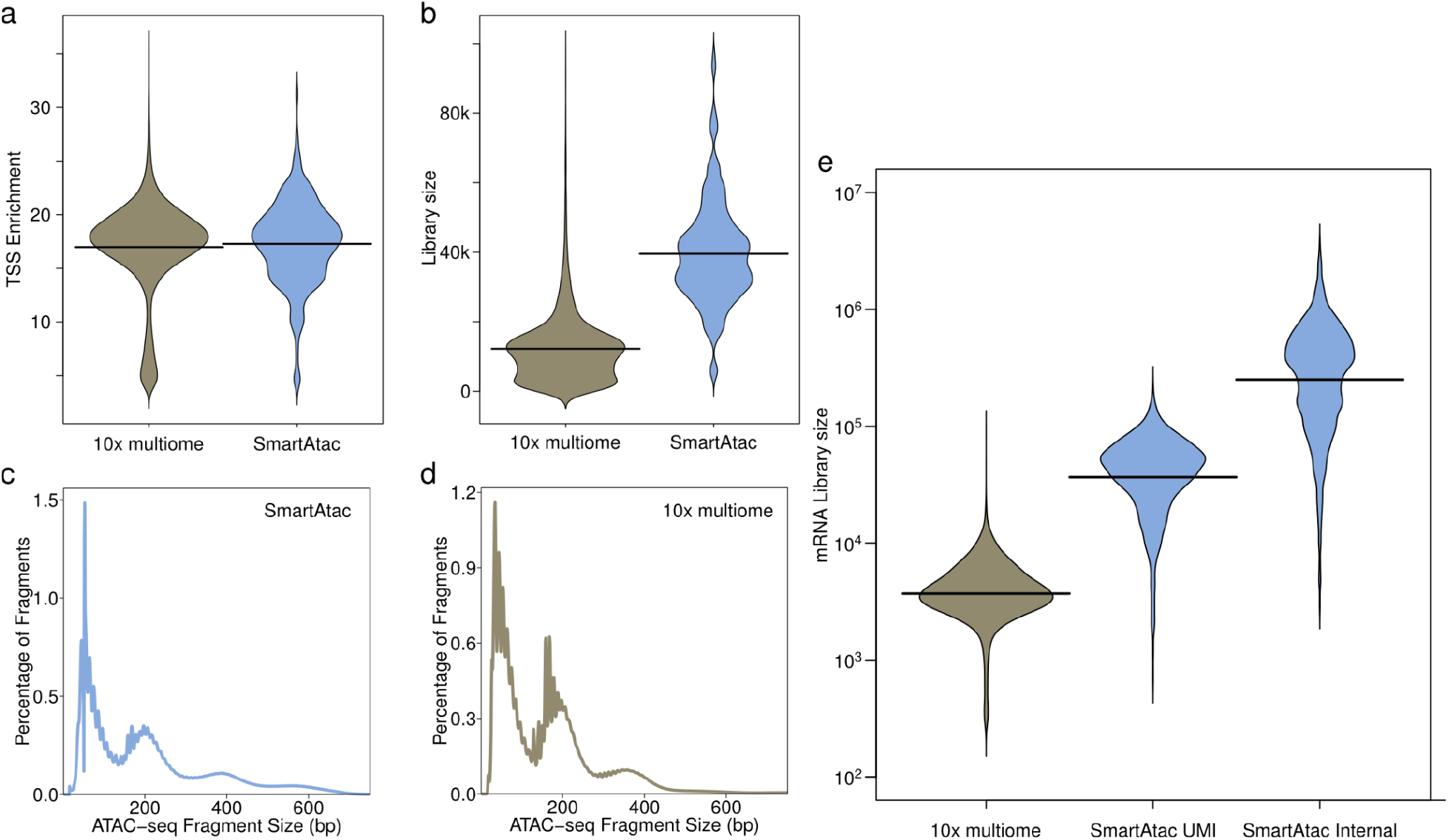
Quality of Smart3-ATAC compared to 10x multiome v1. a) Enrichment of reads around the TSS of annotated genes. b) Sizes of scATAC libraries. c) Fragment sizes for Smart3-ATAC d) Fragment sizes for 10x multiome. e) Sizes of mRNA libraries. 10x multiome (4 302 UMIs/cell average), Smart3-ATAC (46 501 5’ UMI reads/cell and 398 458 internal reads/cell average).

Using the more abundant cytosolic mRNA instead of nuclear mRNA allows us to obtain the highest possible quality transcriptomic measurements. Notably, our mRNA-seq is based on the Smart-seq3, which combines UMI-based absolute quantification of transcript abundance with highly sensitive whole-transcript coverage. 5’ UMI counts for Smart-seq3 are more than 10-fold higher than the 3’ UMI counts from 10x multiomic analysis (Fig 2e), a difference which is most likely primarily due to the fact that we analyze cytosolic mRNA rather than the much less abundant nuclear RNA. Internal non-UMI exon counts are almost a factor of 10 higher, and can be readily increased by deeper sequencing.

An important use of joint transcriptomic/accessibility measurements is to predict specific gene regulatory activity. This is achieved by calculating the correlation or predictive power of the accessibility of a non-coding regulatory element and transcript abundance of a gene. As expected, Smart3-ATAC found a marked enrichment of gene associations to genes in close association to their TSS, with 48% of high confidence (p<1e-20) cis-peaks occurring within 100bp of their associated TSS. However, many genes such as Pbx1 (a pioneering transcription factor important in development of several organs(Kim et al. 2002; Schnabel, Selleri, and Cleary 2003; Sanyal et al. 2007)) completely lacked any correlation between promoter accessibility and transcript abundance, and are instead potentially driven by distal enhancers (Fig 3).

**Figure 3.**
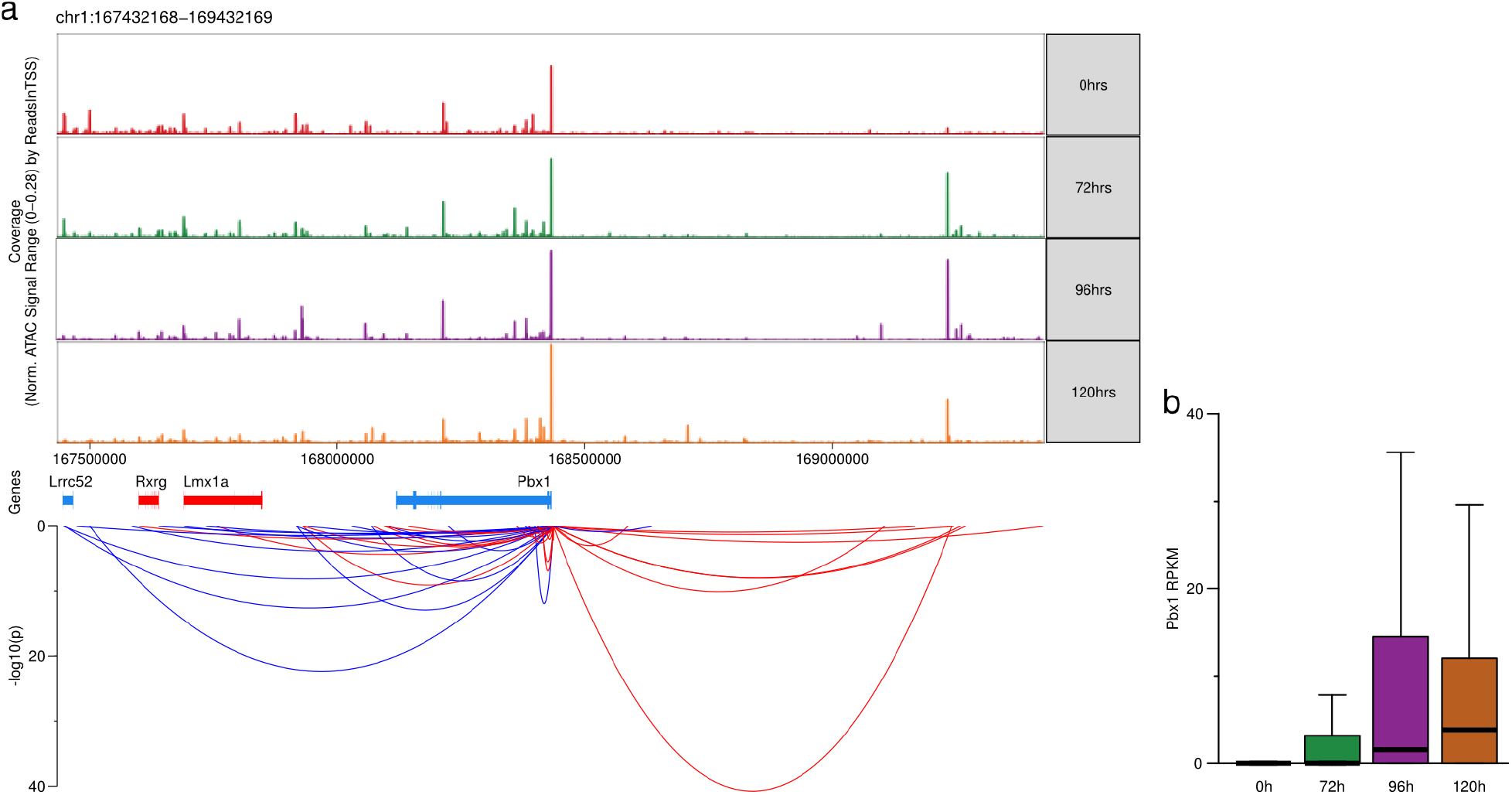
Pbx1 expression is independent of promoter accessibility but highly correlated with distal enhancer. a) Accesibility around Pbx1 and significance of Pbx1 expression. Top: pseudo-bulk tracks divided by time-point. Middle: protein-coding genes in the region, red indicates genes on + strand, blue - strand. Bottom: correlations between Pbx1 and surrounding accessibility peaks in the Gastruloid data set. Ribbons connect Pbx1 TSS and peaks, with the y-axis indicating significance of the correlation. b) Transcription level of Pbx1 across gastrulation time-points.

## Discussion

We describe here Smart3-ATAC, a method for joint mRNA and accessibility measurements from single cells. Contrary to most similar methods, Smart3-ATAC allows obtaining highest-quality data rather than maximizing number of cells assayed. This approach has several advantages. First, using multiwell plates has many benefits over droplets. Since microwells are individually addressable, multi-tiered experiment setups are possible since specific sets of cells can be re-analyzed after the initial analysis. For example, we can re-sequence rare cells to obtain data of higher resolution or we can genotype cDNA of specific cells by qPCR to determine an accurate relative allele-specific expression value for the gene. Second, enhancer/gene association analysis might benefit in a superlinear fashion from high sensitivity measurements. This analysis is dependent on co-occurrence of mRNA reads and scATAC reads in the same cell, which means that increases in yield from both modalities will have an exponential effect on the number of informative cells for a given transcript/enhancer pair. The drawback is increased per-cell cost and decreased throughput, precluding the use of Smart3-ATAC in ultra-large-scale projects. However, for scientific questions where cell number is limited, or where many samples with relatively few cells each have to be analyzed in parallel, high-throughput methods can be more expensive and laborious since while per-cell cost is much lower, the per-experiment cost is typically higher.

## Methods

### scATAC library preparation

Single cells were FACS sorted into 384 well plate that each well contains 3μl lysis buffer (0.03ul 1M Tris-pH7.4, 0.0078μl 5M NaCl, 0.075μl 10% IGEPAL, 0.075μl RNases Inhibitor, 0.075μl 1:1.2M ERCC, 2.7372μl H_2_O). Right after the sorting, the plate underwent centrifuge at 1800g 4°C 5mins, placed on ice 5mins, vortex 3000rpm 3mins, and centrifuge again at 1800g 4°C 5mins to lyse the cells and spin down the nucleus. 2μl of the supernatant was then carefully moved (at <0.5μl/s) to a new 384 well plate for Smart-seq3 mRNA library preparation and the nucleus remained in the original well for scATAC library preparation. The scATAC in situ tagmentation was performed with 2μl of the Tn5 tagmentation mix (0.06μl 1M Tris-pH 8.0, 0.0405μl 1M MgCl_2_, 0.2μl Tn5) and incubating in 37°C for 30mins. Tn5 underwent a His-tag (Dynabeads His-Tag Isolation & Pulldown, Invitrogen, 10103D) clean-up according to the manufacturer’s protocol to remove unbound oligos before using.

After the clean-up, we use the Tn5 assembled oligos to indicate their concentration (Qubit). We applied 4ng/μl as a standard concentration to adjust the use of Tn5. After the tagmentation, 2μl of the supernatant was aspirated and the nucleus were then washed once with 10μl ice-cold washing buffer (0.1μl 1M Tris-pH7.4, 0.02μl 5M NaCl, 0.03μl 1M MgCl_2_, 9.85μl H_2_O). The remaining Tn5 was inactivated by adding 2μl 0.2% SDS-Triton X-100 and incubating at room temperature for 15mins and 55°C for 10mins.

Before barcoding PCR, 2μl Proteinase (0.0107au/mL) was applied to liberate genomic DNA from chromatin by 55°C 1h treatment and followed by 70°C 15mins heat inactivation. Barcoding PCR was done by KAPA HiFi PCR Kit (Roche) in a final 25μl reaction (11.5μl H2O, 5ml 5X reaction buffer, 0.75μl dNTP, 0.5μl KAPA HiFi DNA Polymerase, 2μl barcoding primers). The PCR program was 72°C/15 min, 95°C/45s, [98°C/15 s, 67°C/30 s, 72°C/1 min] x 22 cycles, 72°C/5 min, and then 4°C hold. After the PCR, 2μl of each well was pooled and cleaned-up twice using SPRI-beads (at 1.3X volume) and the library was sequenced on Illumina NextSeq 550 as paired-end, dual index (37+37+8+8).

## Data analysis

mRNA-seq data analysis was performed as in Zachariadis et al(Zachariadis et al. 2020). For the joint scATAC-seq data, raw sequence reads were trimmed based on quality (phred-scaled value of >20) and the presence of illumina adapters, and then aligned to the mm10 genome build using BWA(Li and Durbin 2010). Reads that were not mapped, not primary alignment, missing a mate, mapq <10, or overlapping ENCODEs blacklist regions(Amemiya, Kundaje, and Boyle 2019) were removed. A custom script was used to summarize the paired-end reads into a de-duplicated fragment file suitable for downstream analysis.

scATAC-seq data was analyzed using ArchR. Analysis of accessibility peak to gene association was performed using a linear model implemented in the Matrix eQTL package(Shabalin 2012). First, we called accessibility peaks via ArchR and obtained a binary matrix of peaks-by-cells where each field was 1 if the cell had a read in the peak and 0 otherwise. This matrix was used as alt variants in the analysis. Gene expression values were normalized and log-transformed, and the association analysis was run without covariates and limited to only cis-acting effects (distance between peak and gene < 1E6 bases).

The 10x genomics multiome dataset(“[No Title]” n.d.) was downloaded and loaded directly into ArchR.

